# Heritability and host genomic determinants of switchgrass root-associated microbiota in field sites spanning its natural range

**DOI:** 10.1101/2022.06.09.495345

**Authors:** Joseph A Edwards, Usha Bishnoi Saran, Jason Bonnette, Alice MacQueen, Jun Yin, Tu uyen Nguyen, Jeremy Schmutz, Jane Grimwood, Len A. Pennacchio, Chris Daum, Tijana Glavina del Rio, Felix B. Fritschi, David B. Lowry, Thomas E. Juenger

**Affiliations:** Department of Integrative Biology, University of Texas, Austin; Genome Sequencing Center, HudsonAlpha Institute for Biotechnology; Joint Genome Institute, Lawrence Berkeley National Laboratory; Department of Plant Sciences, University of Missouri; Department of Plant Biology, Michigan State University

## Abstract

A fundamental goal in plant microbiome research is to determine the relative impacts of host and environmental effects on root microbiota composition, particularly how host genotype impacts bacterial community composition. Most studies characterizing the effect of plant genotype on root microbiota undersample host genetic diversity and grow plants outside of their native ranges, making the associations between host and microbes difficult to interpret. Here we characterized the root microbiota of a large population of switchgrass, a North American native C4 bioenergy crop, in three field locations spanning its native range. Our data, composed of >2000 samples, suggest field location is the primary determinant of microbiome composition; however, substantial heritable variation is widespread across bacterial taxa, especially those in the Sphingomonadaceae family. Despite diverse compositions, we find that relatively few highly prevalent bacterial taxa make up the majority of the switchgrass root microbiota, a large fraction of which is shared across sites. Local genotypes preferentially recruit / filter for local microbes, supporting the idea of affinity between local plants and their microbiota. Using genome-wide association, we identified loci impacting the abundance of >400 microbial strains and found an enrichment of genes involved in immune responses, signaling pathways, and secondary metabolism. We found loci associated with over half of the core microbiota (i.e. microbes in >80% of samples) regardless of field location. Finally, we show a genetic relationship between a basal plant immunity pathway and relative abundances of root microbiota. This study brings us closer to harnessing and manipulating beneficial microbial associations via host genetics.

## INTRODUCTION

Recent insight into the composition, ecology, and functional importance of the plant microbiome has greatly increased interest in the potential to harness root microbiota to sustainably increase crop resilience and yield. Microbial inoculants have historically been discussed as a means to achieve this goal, but more recent calls for using plant breeding to enrich beneficial bacteria from the native microbiota have begun to emerge. A roadblock hampering this effort is a lack of understanding about which microbes can respond to breeding practices, whether breeding can instill consistent effects on microbial assemblages across differing environments, and which genes and pathways from the host can be adjusted to modify microbiomes.

Plant root bacterial microbiomes are derived from soil-borne communities, for which membership is largely driven by environmental factors such as geography and climate ^1,2^, land use history ^3^, and seasonal variation ^4–6^. The host plant exerts additional influence over its microbiota through active and passive mechanisms, resulting in filtered subsets of soil microbiota often composed of consistently enriched microbial taxa on and inside root tissue. Given that microbiota can impart positive and negative outcomes on plant health, especially under varying environmental conditions, it follows that the filtering process may be under selection and lead to microbe-mediated local adaptation ^7^.

Heritable variation is required for a trait to respond to selection. Indeed, several recent studies indicate that abundances of rhizosphere and root microbiome members are heritable ^8–13^, i.e. specific microbes and overall community composition vary depending on the genetic background of the host. These studies allude to the possibility of enriching for beneficial microbial associations through breeding, but given that most of these types of studies only look at a few host genotypes and/or grow host plants outside of their native ranges, the role of host genetics in root - microbe interactions has been difficult to interpret. Furthermore, given our relatively recent understanding that features of the microbiome are heritable^14–16^, genomic loci underlying root associated microbiome composition are still largely uncharacterized. There are notable exceptions however: Deng et al used the Sorghum Association Panel to uncover loci impacting rhizosphere community composition ^17^. Bergelson et al. performed GWAS on Arabidopsis root (and leaf) microbiome community metrics including richness and principal coordinates based upon community dissimilarity ^18^. Uncovering the effects of host genetics on microbiomes across multiple native environments remains incomplete, but these studies provide exciting avenues to leverage host genetics to enrich for beneficial properties of the microbiome.

Switchgrass (*Panicum virgatum*) is a wild C4 perennial prairie grass native to North America and has been championed by the US DOE as a potential biofuel crop due to its biomass yield potential when grown in marginal soil with minimal agricultural inputs. Its interesting biological features and important environmental and economic impact have made switchgrass a popular model to investigate root-associated microbiota assembly, especially in the rhizosphere <u>(Singer et al. 2019; Ulbrich et al. 2021)</u>. Most recently, Sutherland et al. used a panel of switchgrass genotypes grown in a single location in the northeast United States to uncover the role of host genotype on rhizosphere bacterial assemblages ^21^. The authors of this study used GWAS to uncover putative loci affecting the abundance of several bacterial families in the rhizosphere and found gene ontology enrichments for diverse sets of functions. Still, relatively little is known about how host genetics drive tightly adhering / endophytic root-associated bacterial communities.

In this study we addressed the following questions: 1) What bacteria are prominent members of the switchgrass root-associated microbiome when plants are grown across their natural range? 2) How does the effect of host genotype compare to that of the environment when determining the composition of root-associated bacterial microbiota? 3) Which microbial lineages show heritable variation in roots, and is heritability consistent across field sites? 4) Which host genomic loci impact the abundance of root associated bacteria? 5) Does microbial abundance show patterns of association with host immunity variation. Answering these questions will bring us closer to harnessing and manipulating beneficial microbial associations via host genetics.

## RESULTS

### Field site is a primary determinant of switchgrass root microbiota composition

We used a population of fully resequenced switchgrass (*Panicum virgatum*) natural accessions that were clonally replicated and grown in field sites at Austin, TX, Columbia, MO; and Kellogg Biological Research Station, MI (from here on referred to as ATX, CMO, and KMI, respectively Fig 1A, map inset) to uncover the role of environmental variation and host genetics in shaping root microbiota composition. These plants had been established for two years, show signatures of local adaptation ^22,23^, and have served as an important resource for switchgrass researchers. We first investigated the effect of field site on root bacterial microbiota. Principal coordinate analysis (PCoA) revealed three dominant clusters which were location-specific (Fig. 1A) and the significance of this observation was confirmed using perMANOVA (R2 = 0.51, P <0.001). While the communities showed large differences between field sites at the amplicon sequence variant (ASV) level, we found that phylum level relative abundances were largely consistent between sites (Fig. 1B). Actinobacteria and Proteobacteria (namely Alpha and Gamma-proteobacteria) were dominant phyla associated with switchgrass roots at every site, which is consistent with most other terrestrial, non-flooded, plant microbiota studies.

**Figure 1.**
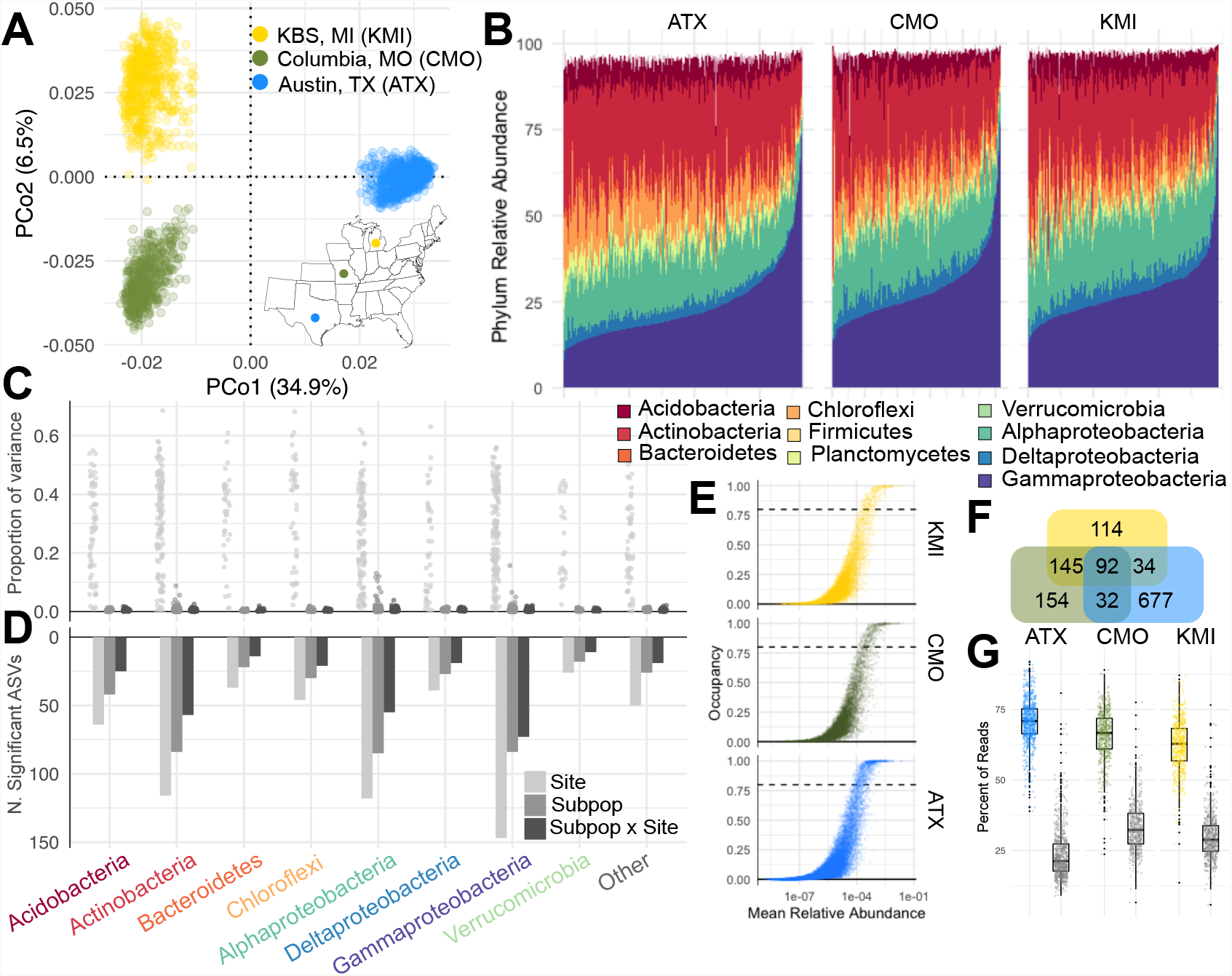
Field site is the primary determinant of switchgrass root microbiota composition. A) Principal coordinate analysis based on Bray-Curtis dissimilarities. Inset: map of field locations, colors match those in the figure legend. B) Relative abundance of phyla and Proteobacterial classes in every sample at each site. C) Effect sizes for Site, Host Subpopulation, and Subpopulation x Site for ASVs in dataset broken down by phylum / class. D) Number of ASVs with significant contrasts from the models displayed in panel C. E) Prevalence / abundance curves for each field site. Each point represents a single ASV and the black dashed line is the 80% prevalence threshold used to call core taxa. F) Venn diagram displaying overlaps of core microbiota from each site. G) Fraction of reads belonging to the core microbiota at each site (colored boxes) and the shared core microbiota (92 overlapping microbes from panel F, gray boxes).

A recent population genomic study of switchgrass found that tetraploid switchgrass can be broadly classified into three genetic subpopulations: Gulf, Midwest, and Atlantic ^22^. The ranges for these subpopulations are largely distinct (See Fig. 2A), with Gulf occupying habitats in the southern US, Atlantic occupying the Atlantic seaboard, and Midwest spread across northern latitudes. We compared the effect of field site, host subpopulation, and their interaction using linear models run on bacteria present in ≥ 50% of the samples study-wide. The effect of field site was much larger than the secondary effects of host subpopulation and subpopulation x site interactions (Fig. 1C). We then compared the variance explained by site between bacterial phyla / classes to better understand how experimental factors impact broader taxonomic groupings. Effect sizes were largely consistent between these groups, with the exception of Chloroflexi and Actinobacteria, which showed larger effect sizes than Deltaproteobacteria (P < 0.05, Tukey’s Post-hoc Test). The large influence of field site on ASV relative abundance was also visible in the number of ASVs which exhibited significant differences in relative abundance across field sites (Fig. 1D).

**Figure 2.**
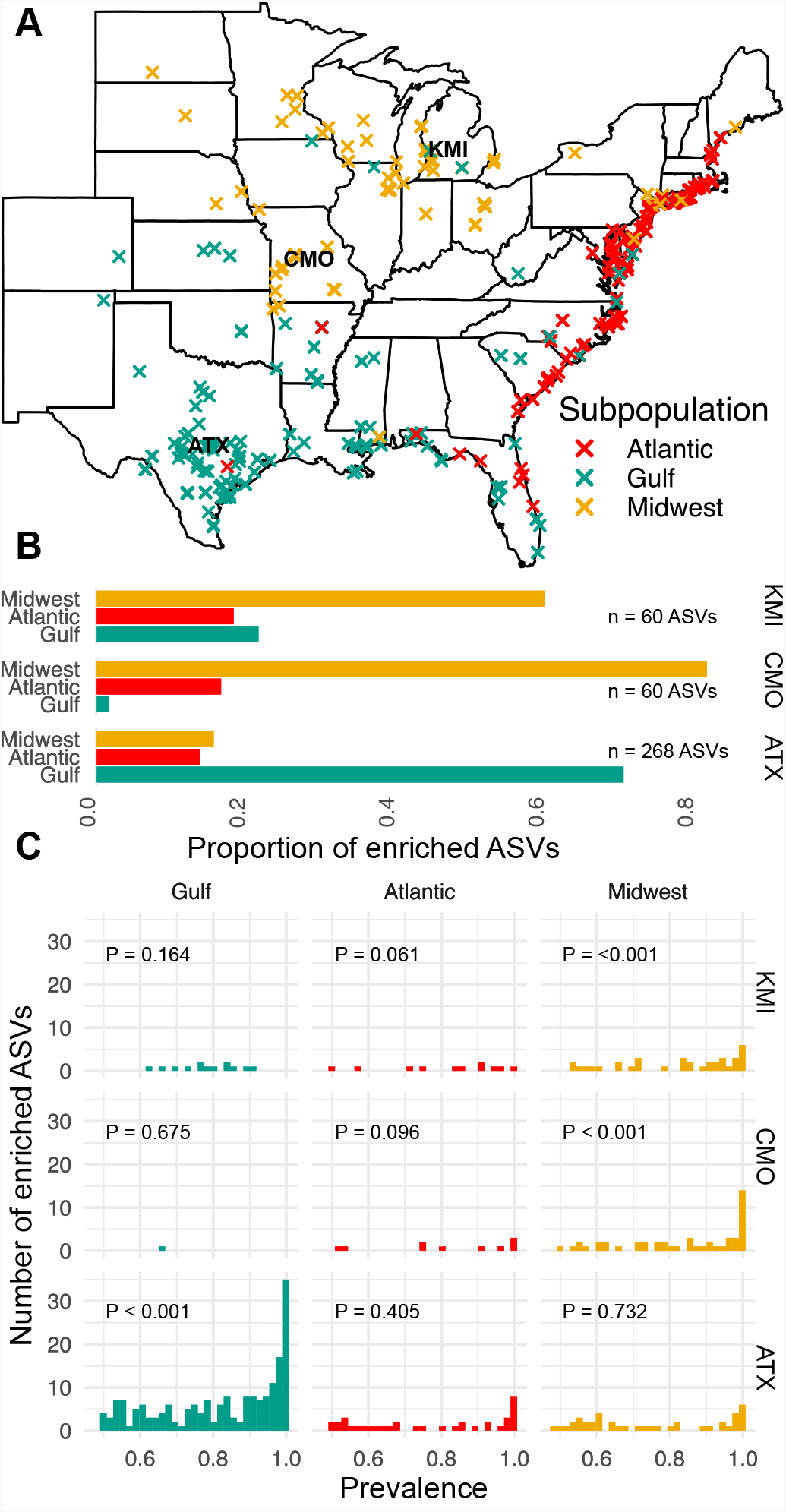
Plants show evidence of affinity to local bacterial strains. A) Map depicting locations where individuals within the population were collected. Colors represent their subpopulation memberships. Field sites are depicted with their three letter abbreviations. ATX = Austin, TX; CMO = Columbia, MO; KMI = KBS, MI. B) Proportion of ASVs showing specific enrichments in one subpopulation compared to the other two broken up by site. C) Histograms of microbial prevalence showing specific enrichments by subpopulation and site. P values represent the significance of the mean prevalence being greater than that of the background distribution. This was calculated by randomly drawing the number of enriched ASVs from the background distribution and asking how often we saw a mean prevalence greater than that of the focal set.

We next evaluated the relationship between ASV occupancy and mean relative abundance at each site (Fig 1E). Our study used an atypically high depth of sequencing (Supp. Fig. 1) which gave us greater confidence in assessing presence / absence of microbes in samples. In general, we found that ASVs with greater relative abundances were also present in a higher proportion of root microbiomes. We next defined site-specific core microbiota; to be consistent with other studies, we used a threshold of 80% occupancy ^8^ (Supp. Table 1). ATX had the most ASVs passing this occupancy threshold (Fig. 1F); we expected this, because we sequenced ATX samples at greater sequencing depths than the other two sites (Sup. Fig. 1, See Methods). Still, we found that each site hosted overlapping core microbiota: For all three sites, an overlap of 92 core microbes was found. CMO and KMI shared the most ASVs. The site-specific core microbiota typically comprised ∼60-70% of the total microbial population (Fig 1G, colored boxplots) within each respective site, while the shared core microbiota made up ∼25% of the total population (Fig 1G, gray boxplots). Thus, though field site acts as the primary determinant of switchgrass root-associated microbiota composition, large proportions of switchgrass root assemblages are shared between locations as a set of core microbes.

### Evidence of affinity between host genotypes and local microbiota

Our analyses revealed that host subpopulation and subpopulation by location interactions are important determinants of microbiota composition (Fig. 1C and D). Because the three switchgrass subpopulations are largely constrained to distinct geographic regions (Fig. 2A), we hypothesized that plants grown closer to their native habitat would show affinity for the microbes that persist and are abundant within these ranges. If this was true, then we would expect, at each site, that more ASVs would show preferential colonization of individuals in the subpopulation grown in its native range than in the other two subpopulations. To test this, we used linear models to analyze the abundance of ASVs within each site and contrasted the abundances between the different subpopulations. We defined a specific association as occurring if the relative abundance of an ASV was significantly greater in one subpopulation compared to the other two. Gulf plants in their native ATX site had the most specific associations, while Midwest plants enriched the most ASVs in native CMO and KMI sites (Fig. 2B, Supp. Table 2), supporting the notion that subpopulations enrich more microbes in their native habitats. Furthermore, we found the ASVs with subpopulation specific associations also tended to have significantly greater prevalence (Fig. 2C), but only for subpopulations growing within their native range. That is, ASVs with specific associations in the Gulf subpopulation had significantly greater prevalence than the background distribution at the ATX site, but not the other two sites. Likewise, microbes with specific associations in the Midwest subpopulation showed significantly greater prevalence in both CMO and KMI sites compared to the background prevalence distributions (Fig 2C). These comparisons suggest there is preferential sorting of local microbiota onto locally adapted plant genotypes, especially for highly prevalent microbes.

### Switchgrass root microbiota show widespread heritable variation and genotype by environment interactions

Our analysis of switchgrass subpopulation effects on microbiota abundances underscores the importance of broad level host genotype in modulating root microbiome assembly. We next used an approach which incorporates a kinship matrix denoting finer genetic relationships among individuals of the population into the model to estimate how host genetic variation contributes to variation in microbe abundance. We used a suite of mixed effects models to partition additive genetic variance in microbial abundance (V_A_) using the host population genetic relationship matrix and how V_A_ differs across the three environments (V_GxE_) with a compound symmetry model. Because microbiomes can be defined and analyzed at various taxonomic levels by aggregating counts at nodes of the bacterial phylogenetic tree, we tested the affect of host genotype on the relative abundance of taxa at various taxonomic levels. Across each taxonomic level both V_GxE_ and V_A_ significantly explained variation in microbial abundance (Fig 3A, Supp. Table 3). For microbial features within the top 10th percentile for V_A_ and V_GxE_, we found generally increasing estimates for V_A_ and decreasing estimates for V_GxE_ from phylum to ASV (Fig. 3B). We next asked whether taxonomic groupings of microbes at the ASV level were more likely to be under the influence of host genetics. Significant, non-zero V_A_ and V_GxE_ were widespread across the microbial phylogeny, however specific orders were overrepresented in the data (Fig. 3C). In particular each tested ASV within the orders Sphingomonadales, Subgroup 6 (Acidobacteria), Gammaproteobacteria Incertae Sedis displayed significant V_A_ or V_GxE_. We next compared the contribution of V_A_ to V_GxE_. In general, we found that more microbial features showed greater V_GxE_ and this was consistent across taxonomic levels (Fig. 3D). The prominence of GxE suggested that levels of V_A_ differ between locations. To better understand the contribution of V_A_ within each site, we fit an unstructured model to ASVs which allowed for site-specific V_A_ and as many unique covariances as site combinations. We applied these models to ASVs with prevalences > 80% in at least two field sites (Fig. 3E), finding similar trends to the compound symmetry model (Supp. Fig. 2). When analyzing the core microbiota (i.e. the 92 ASVs with prevalence >80% in all three sites), we found 95 instances of significant site-specific V_A_ spread across 64 unique ASVs (Supp. Table 4). CMO had the most ASVs displaying significant V_A_ (n = 38) while KMI had the least (n = 24). We also tested if there was a genetic association between the abundance of an ASV across multiple sites by focusing on the genetic covariance of root-associated microbial traits across sites. Genetic covariances were mainly positive (Supp. Fig. 3A) and site comparison had a significant effect on covariance strength (P = 0.005, ANOVA). Specifically, we found that CMO/KMI covariances were significantly greater than those from ATX/KMI (adjusted P = 0.006, Tukey’s Post Hoc Test), but not ATX/CMO (P > 0.05, Tukey’s Post Hoc Test). We tested for ASVs that showed significant genetic covariance between sites and found 78 total significant estimates spread across 59 unique ASVs. Similar to the aggregate genetic covariance distributions, we found the most cases of significant genetic covariance between CMO/KMI, while CMO/ATX and KMO/ATX had equal instances of significant estimates (Sup. Fig. 3B). Together, these results indicate the host genetics plays a significant role in modulating an extensive phylogenetic swath of root-associated microbiota, that some bacterial clades are more likely to display heritable variation, and that genotype by environment interactions are widespread determinants of bacterial relative abundances on switchgrass roots.

**Figure 3.**
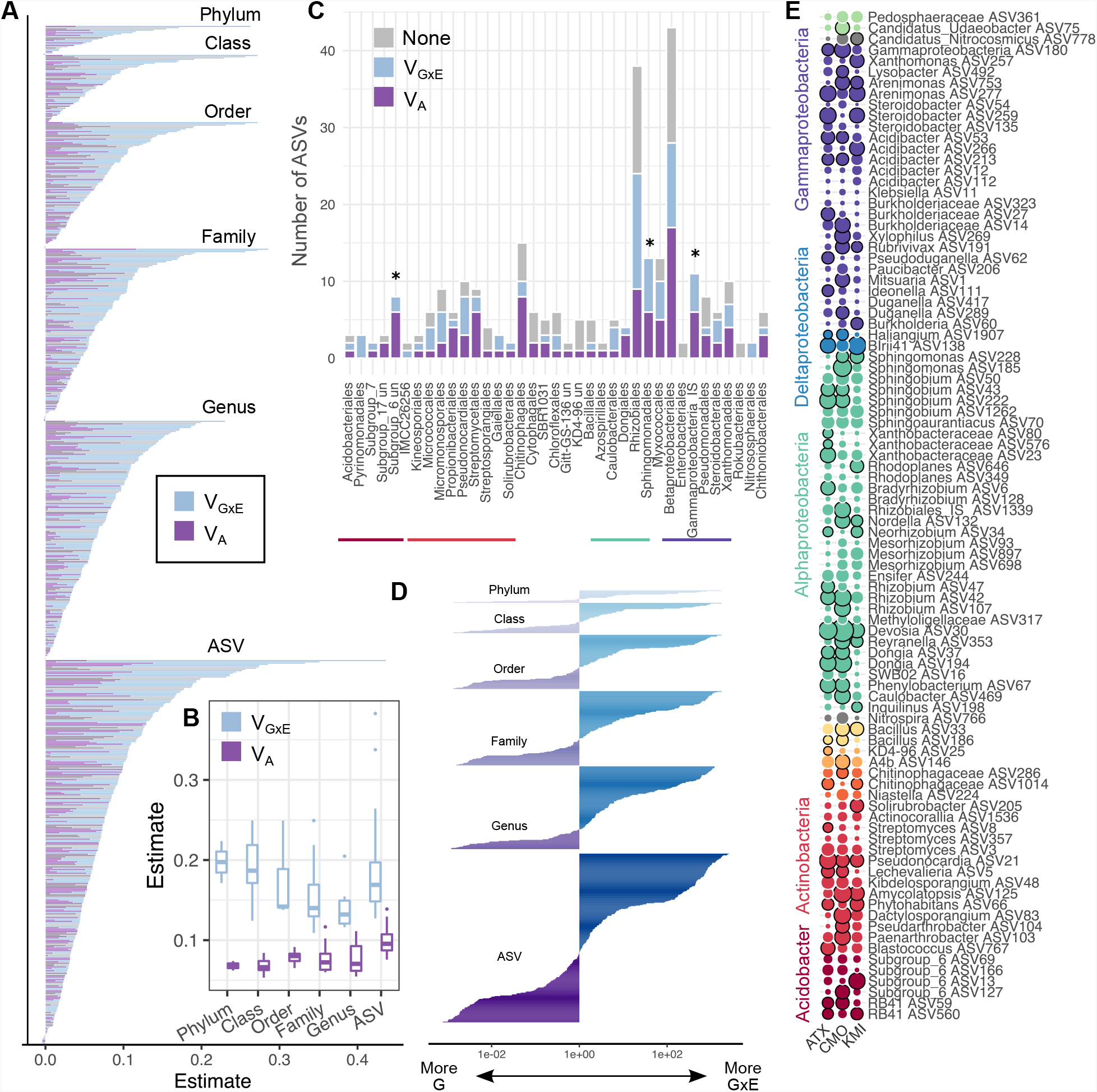
Switchgrass root microbiota show widespread heritability which is influenced by field site differences. A) Variance components for aggregated abundances of different taxonomic levels and for ASVs. To be included in the models, features must have been present in greater than 80% of the samples, study-wide. B) The relationship between genetic variance components and microbial taxonomic rank C) The number of ASVs showing either significant GxE, VA, or no association to host genotype D) Comparison of the magnitude of VA vs GxE is presented as the log fold-change in the ratio of VA to GxE for measured units within each taxonomic level. E) VA estimates for the core microbiota present at every site. The size of the circles indicate the magnitude of estimated VA and dark perimeters of the circles indicate a significant association (FDR < 0.1).

### GWAS reveals microbiota assembly is a complex trait with extensive pleiotropy

After establishing that host genotypic variation influences the abundance of bacterial taxa, especially within single field sites, we next asked if host genomic regions responsible for heritable variation in associated bacteria could be localized with a genome wide association study (GWAS) framework. We first performed GWAS on community composition using the first three principal coordinates for each site (Supp. Fig. 4). Significant associations between SNPs and community composition were detected for each site, albeit on different PCo axes. These results indicate that variation in community composition is associated with host allelic variation. To better understand how host allelic variation influences individual microbes, we extended our analysis to perform GWAS on each ASV x site combination. We analyzed ASVs present in at least 80% of the samples, resulting in 1019 independent analyses of ASV x Site combinations. GWAS results were examined using a genome-wide significance threshold of 5×10^−8^ to identify SNPs associated with the abundance of various microbes, a common cutoff used in microbiome GWAS studies where many phenotypes are analyzed together ^24,25^. Using this criterion, we found 1,153 SNPs associated with 459 ASV x Site combinations. Most ASVs with significant SNP associations were from the ATX site (253 ASVs), while CMO and KMI had similar numbers of ASVs with associated SNPs (101 and 105 ASVS, respectively). Taxa with associated SNPs were diverse, but no bacterial orders were over-represented (Fig 4A-C). Most ASVs with associated SNPs were specific to field sites; however, of the 179 ASVs that were tested in multiple sites, 50 showed associations across multiple field sites, with 9 showing associations across all three sites (Supp. Fig. 5D). In line with our heritability analysis, bacteria within Sphingomonadaceae featured prominently among ASVs with GWAS hits across multiple sites: 7 of the 10 ASVs within this family showed hits across 2 or more sites and 2 *Sphingobium* ASVs had at least one significantly associated SNP at all three sites (Fig. 5D).

**Figure 4.**
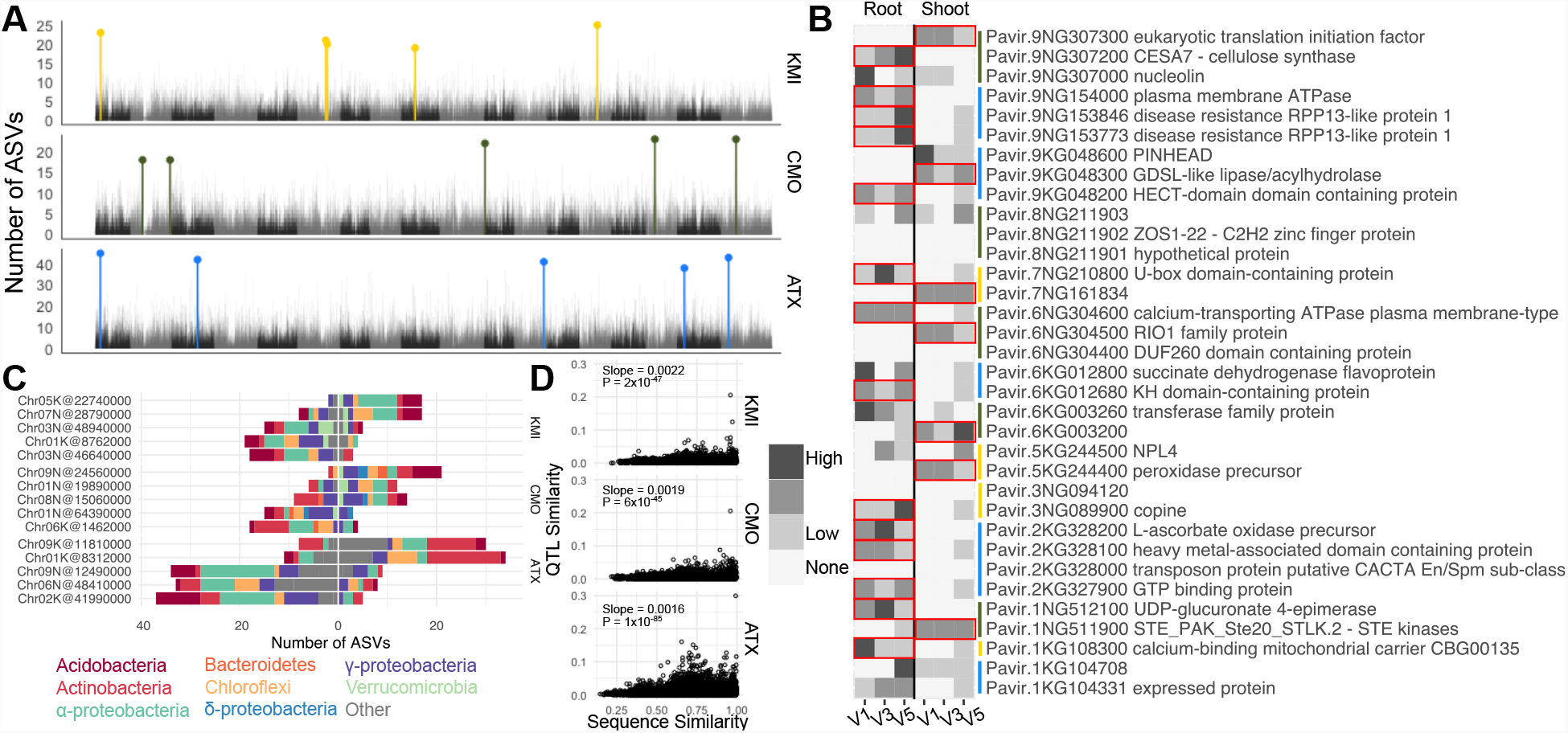
Pleiotropic loci influencing root microbiota. A) Number of ASVs detected in the 0.5% tails of the ASV x site GWAS p-value distributions. The top 5 most frequently observed genomic bins for each site are highlighted in site-specific colors. B) Candidate genes underlying the pleiotropic loci and their expression pattern in switchgrass roots and shoots. V1-V3 represent phenological stages of the plant and red boxes around expression values represent genes differentially expressed between roots and shoots (FDR < 0.05) C) Taxonomic breakdown of ASVs affected by putatively pleiotropic loci. D) Comparison of QTL similarity (1 -Jaccard Dissimilarity) and ASV sequence similarity.

**Figure 5.**
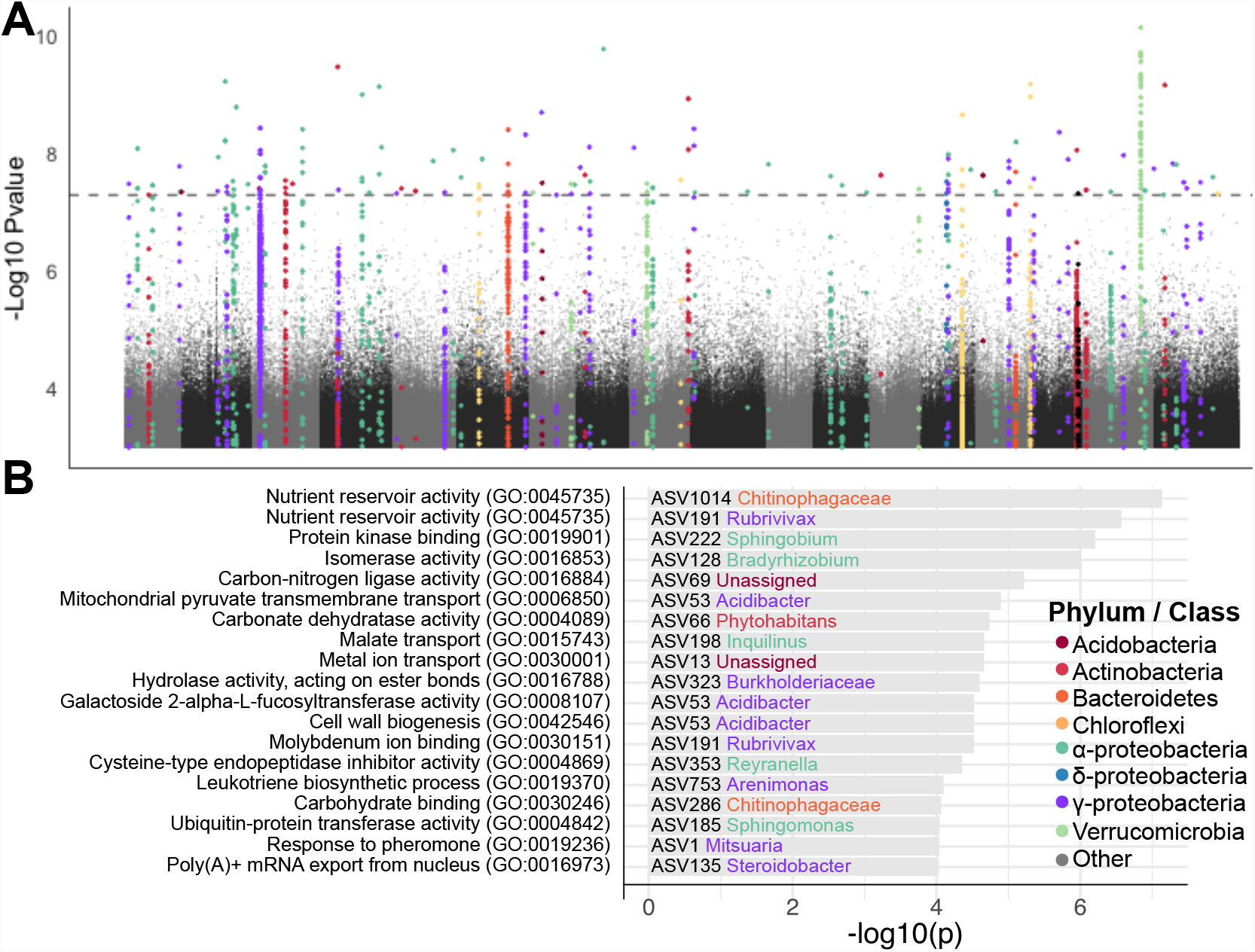
GWAS reveals loci associated with core switchgrass root microbiota. A) Manhattan plot showing the association between SNPs and abundances of core ASVs. P values are derived from combining P-values using Fisher’s method. Peaks are colored by the Phylum / Class of the ASV. B) The most strongly enriched Gene Ontology (GO) terms within the core ASV GWAS tails.

We next asked whether any host genomic loci affected multiple microbial taxa (i.e. had statistically pleiotropic effects on microbiota and from here on referred to as pleiotropic loci) by compiling the 0.5% tail of 25 kB genomic bins into a quantitative trait locus (QTL) x ASV matrix for each site (see Methods). We first investigated the most commonly observed 25 kb genomic bins for each site by selecting the top 5 loci associated with the most ASVs within each site (ATX = 38-45 ASVs; CMO = 18-23 ASVs; KMI = 19-25 ASVs, Supp. Table 5). Two pleiotropic loci overlapped with loci detected from our initial GWAS on community metrics (Supp. Fig. 4; CMO:Chr01N and ATX:Chr02K), indicating that while some pleiotropic loci account for larger trends in community composition, most identify variation not seen along the first three axes of community composition.

To better characterize the candidate genes underlying these loci, we next compiled expression patterns for genes within these intervals. Most loci contained genes displaying higher expression patterns in switchgrass roots than shoots, implicating promising candidate genes affecting multiple microbiota members. These included several proteins involved in calcium signaling, immunity, and secondary cell wall biosynthesis. The microbes associated with pleiotropic loci were taxonomically diverse, with multiple bacterial phyla affected by each locus. In general, the additive effects of the QTL were largely consistent in sign across the different ASVs. This observation was also reflected in the taxa being affected by the loci: several loci show patterns where the relative abundances of Actinobacteria, Chloroflexi, or Alphaproteobacteria ASVs had consistent effect signs. This observation led us to the hypothesis that there may be an association between the QTL landscape and phylogenetic relationship for pairs of microbes. We found a positive and significant association between the sequence similarity of ASVs and their associated QTLs. This association differed weakly but significantly between sites with ATX showing a weaker correlation than CMO or KMI (P = 0.06 and 0.0015, respectively). Each site had a closely related ASV pair which stood out in terms of shared QTLs. These included two *Sphingobium* ASVs in ATX, *Bacillus* in CMO, and *Acidibacter* in KMI. Together these results indicate that host genomic variation can have pleiotropic effects on microbiota and that the abundances of related microbes are more likely to be affected by the same host loci.

The pleiotropic loci included several promising candidate genes, but to have a more robust understanding of the functional categories influencing switchgrass root associated microbiota we performed gene ontology (GO) enrichments for annotated genes underlying the ASV x QTL matrix. We found that 789 of the ASV x site combinations displayed at least one significant GO enrichment. The most commonly observed GO term enrichments showed overlapping as well as contrasting patterns between sites (Supp. Fig. 6, Supp. Table 6). For example, the terms ‘response to biotic stimulus’, ‘response to auxin’, ‘negative regulation of growth’, and ‘sucrose biosynthesis’ were observed in multiple ASVs across every site, while ‘Defense response’, ‘prophenate biosynthetic process’, and ‘carbohydrate binding’ showed more site-specific patterns. These results indicate that variation in host molecular pathways can influence the abundance of microbiota members and that some pathways are putatively dependent on environmental conditions.

**Figure 6.**
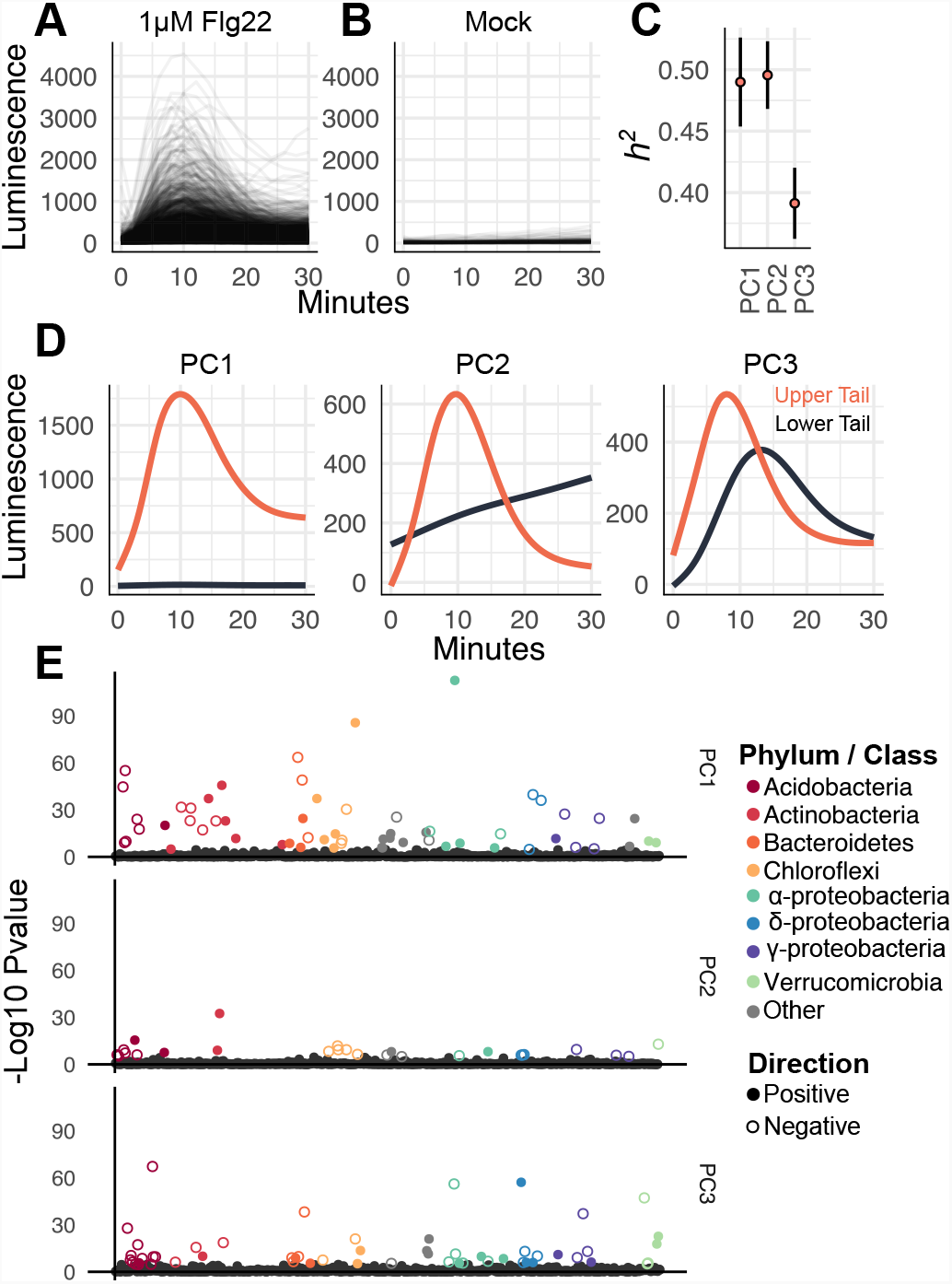
ASV abundances genetically co-vary with pattern triggered immune responses. A) Response curves for the switchgrass population planted at the ATX site for treatment with 1 uM Flg22. B) Response curves for mock inoculated plants. C) Narrow sense heritability estimates for the three PC axes of PTI response variation. D) Smoothed 5% and 95% percent tails of the first three PC axes of PTI response variation. E) Microbial Manhattan plot displaying the p-values for the covariances between ASV relative abundance and the PC axes of PTI variation. Colored circles represent ASVs passing a Bonferroni threshold of 0.05.

To better understand the contribution of loci independent of field site, we subsetted our scans to ASVs which had been tested in every site (i.e. the core microbiota), joining P-values generated during GWAS for a single ASV across each field site using Fisher’s method, a practice commonly used in meta-analyses to identify statistical tests with repeatable signal across multiple trials. A total of 239 SNPs passed a P value threshold of 5×10^−8^, revealing 44 out of 92 core ASVs had a significant association (Fig. 5A, Sup. Fig. 5D). More than half of the ASVs with significant associations (23/44) showed significant GWAS hits across multiple sites (Supp. Fig. 5D and Fig. 5A). Interestingly, some ASVs with combined P-values passing this genome-wide threshold did not display any significant associations in the ASV x site GWAS analyses. For example, *ASV6*, a highly abundant *Bradyrhozobium* strain displayed two significant peaks when P-values were combined that were not present during the initial site by ASV GWAS (Supp. Fig. 5D). These results indicate that leveraging multi-site GWAS by combining P-values can identify loci impacting core microbiota.

We explored the functional enrichments of combined p-value GWAS scans from the core microbiota (Fig 5B, Supp. Table 7, Supp. Table 8). We identified 76 distinct GO terms enriched across 48 core ASVs, some of which have *a priori* implications in microbiome assembly. For example, malate transport and cell wall biogenesis were among the most frequent enriched terms. Malate is a prominent root exudate involved in shaping rhizospheric microbiome composition ^26^ and cell walls form physical barriers as well as energy sources for microbes ^27^. Together this analysis revealed that while observations of loci associated with the abundance of various microbes is environmentally dependent, some loci can be implicated across multiple environments and the processes by which the host plant modulates core microbiota are diverse.

### Pattern triggered immunity responses genetically co-vary with root-associated microbiome composition

Plants surveil their biotic environment through perception of microbial associated molecular patterns, eliciting the activation of the pattern triggered immunity (PTI) pathway. We hypothesized that loci responsible for observed variation in PTI may overlap with host genetic variation controlling microbial abundance. To test this hypothesis we treated leaf disks from the population of plants growing in Austin, TX with Flg22, perhaps the most well studied MAMP. We measured the release of reactive oxygen species (ROS) over time using a well-characterized assay (see Methods). Flg22 elicited a range of ROS burst profiles in the population while mock treated samples did not display the typical response curve of treated plants (Fig 6A). We converted the time series into principal components to better understand the different modes of variation displayed across treated samples. The tails of the PC axes were informative of the type of variation observed in the population (Fig. 6B): PC1 best explained the magnitude of response; PC2 separated plants with acute vs gradual responses; and PC3 showed a timing difference of peak ROS burst. All three axes showed significant *h*^*2*^ ranging from 0.48 to 0.38 (Fig. 6C). These results indicate that switchgrass genotypes significantly vary in their response to the PTI elicitor flg22.

The plant immune system has been implicated to actively shape the microbiome ^28^, therefore we hypothesized that genetic variation for PTI responses may genetically co-vary with abundance of various root-associated microbiota. To test this hypothesis we calculated the genetic co-variances for the PTI PC axes against the relative abundance of core bacterial ASVs in the ATX site. We found significant genetic co-variances across each PTI axis: in total 126 / 739 ASVs showed significant genetic covariances with PTI axes (Bonferroni P < 0.05, Fig 6D). PTI PC1 had the most associations and PC2 had the least. PTI PCs 2 and 3 predominantly had negative co-variances with ASVs while PC1 had a similar amount of positive and negative co-variances. These results indicate that bacterial microbiota show positive and negative genetic correlations with PTI responsiveness and that associations between these traits are not phylogenetically limited.

## DISCUSSION

Here we have used natural switchgrass accessions growing in field sites spanning its native range to evaluate the contribution of environment and host genotype on root-associated bacterial assemblages. Field site was a major determinant of bacterial community assemblages in our study. Within sites, however, host genetics influenced the assembly of bacterial microbiomes, with local microbes preferentially colonizing native genotypes. We found numerous associations between bacterial relative abundances and host genomic loci through a GWAS framework, linking the abundance of taxa to host ontology groups and candidate genes. Our meta-analyses of GWAS scans performed on core ASVs implicated host loci affecting microbiota assembly independent of field location. Finally, we present evidence of correlation between pattern triggered immunity in the host and abundance of bacterial taxa associated with the roots.

### Genotype by environment interactions in host-associated microbiomes

A key finding of our study was that relative abundances of bacteria were strongly influenced by the interaction of host genetic variation and field site (Fig. 2 and Fig 3). Further, we found that there were affinities between genotypes growing in their home environments and the local microbiota (Fig 2B). Interestingly, microbes with specific enrichments to local genotypes consistently had higher prevalence than expected (Fig 2C). A potential explanation is that home genotypes, as opposed to foreign genotypes, are more in sync with their native climates, photoperiods, and soil properties. This in turn, may reduce host stress and culminate in the acquisition of consistent microbiota. Alternatively, these results could be explained by a co-evolutionary framework, where evolution in the microbes drives selection on the host, and consequent selection in the microbes ^15^. However, given the stochastic dispersal of soil microbes ^29^, the more likely explanation is one-sided evolution where the local microbe population imposes selection and evolution on the host, rather than the host imposing selection on the microbes. Perhaps the elevated prevalence of enriched microbes equate to more chances for interaction and act to exert stronger selection on hosts (Fig 2C). Another display of GxE was that ASVs rarely showed heritable variation across every site. While GxE for microbial community composition is often complex in these types of studies, the fundamental ‘disease triangle’ framework from the plant pathology field is useful when considering host-microbe associations, regardless of pathogenesis. This theory dictates that for disease to occur, a susceptible host genotype, virulent pathogen, and favorable environmental condition must co-exist. Each of the three points of the triangle can be explored further to explain GxE in root microbiota assemblages. We discuss these three points in the context of our study below.

Firstly, environmental variation occurs in biotic and abiotic flavors, which are not mutually exclusive. Our results indicate that the environment greatly influences the composition of root microbiota at each field site (Fig 1A). Field site had an almost universal effect on the abundance of ASVs (Fig. 1C). The three field sites do differ in their field uses, a factor which can contribute to soil microbiome variation ^3^. Columbia, MO and Kellogg Biological Research Station, MI sites are converted prairie and forest, respectively, and have histories of cultivating crops either agriculturally or experimentally. The ATX field site is located within city limits on a campus with no known history of agricultural cultivation. These land use history differences may explain the relatively large microbiome compositional variation between ATX and CMO / KMI sites. Furthermore, climate patterns are distinct between the sites, CMO and KMI having more similar climate patterns. Alternative favorable conditions may promote growth of certain taxa, which may ultimately influence the abundance of other microbes.

The microbial component of the disease triangle states that a virulent form of the pathogen must be present to infect a host and initiate disease. Implicit to this point is that genetic variation exists for microbes in addition to hosts. Unfortunately, we could not examine genetic variation of individual ASVs in our study, as we based the detection and abundance of different taxa on a small 250 bp segment of a single gene. While this may suffice to classify most microbes down to the genus or species level, it is insufficient to explore bacterial strain level variation. Every ASV in a site is under selective pressure by the local environment. Therefore, an ASV detected at one site will most likely have distinct polymorphisms with adaptive consequences compared to the same ASV at a different field site. Even within sites, ASVs can be composed of multiple microbial lineages, each conveying distinct phenotypes to the host ^30^. Polymorphisms, especially between sites, may preclude the microbe from falling under the genetic influence of the host, explaining why we detect significant heritability for the same ASV in some sites but not others. Nevertheless, we identified ASVs where combined p-values generated from site-specific GWAS helped to uncover loci consistently associated with their abundance. This was the case for half of the ASVs tested under this framework, suggesting that modulation of ASVs through shared mechanisms across field sites is relatively common, yet may not have effects passing a threshold in single ASV x site GWAS. A potential method to study GxE with host associated microbiomes is through construction of synthetic communities, which offer an ecologically relevant, yet controlled system for plants and microbes to interact while experiencing an experimental environment change. However, it must be noted that synthetic communities will remain incomplete representations of root-associated bacterial communities until highly prevalent and abundant, yet recalcitrant microbes become more easily cultivable. For example, strains belonging to Chloroflexi, Acidobacteria, and Verrucomicrobia are prominent members of plant microbial communities, but remain conspicuously absent from root bacterial culture collections ^31–33^.

Finally, the third point of the disease triangle stipulates that a host plant must be susceptible to infection in order for pathogenesis to occur. In our case, this equates to host genetic variants being compatible for colonization by the local ASV. Susceptibility / compatibility, is likely dependent upon both biotic and abiotic environmental conditions. That is, habitat variation and microbial community variation between sites may activate or repress the expression of allelic variants responsible for regulation of microbial colonization. For example, increased temperature attenuates effector triggered immunity in *Arabidopsis*, increasing susceptibility to *Pseudomonas syringae* ^*34*^. Xin et al demonstrate that elevated humidity can greatly influence the pathogenesis of *Pseudomonas syringae*, but in a host genotype dependent manner ^35^. In addition, given that the microbiomes vary substantially between sites, the biotic component of the environment may contribute to expression differences between allelic variants, thus leading to differential enrichment of metabolic, immunity, and developmental pathways. One fascinating angle recently put forward is that microbes which subvert plant immunity may ultimately serve as keystone taxa ^36–38^ by dampening the immune response, allowing other microbiota to side-step the host immune system. Given that the biotic environment largely varies between sites, contrasting keystone taxa may exert alternative effects on different genotypes.

In all of these scenarios it is important to acknowledge that both microbes and plants are sensitive to environmental conditions. Microbes are a critical part of the host plant’s environment, and likewise, the host plant is an environment for the microbes. Environmental variation may change local microbiota community structure which in turn may affect the expression of host genes impacting microbiota assembly.

### Which taxonomic level is appropriate for calculating heritability of bacteria

We find that heritability of microbiota features can be observed across every taxonomic level. Several studies have calculated heritability of rhizosphere or root associated bacteria ^8–10,21^. Typically, the analysis is conducted at the OTU or ASV level (i.e. the taxonomic level with the highest resolution). In the case of Sutherland et al., the authors chose to calculate heritability for aggregated counts of bacterial families. This begs the question: which taxonomic level is appropriate for calculating heritability of host-associated bacteria? Our results indicate that, while individual ASVs displayed the greatest h^2^ on average, relatively high h^2^ can be observed even at the bacterial order and family level. This observation lends some support to the idea that plants do not select for particular microbes (i.e. specific ASVs), but rather for microbes with particular functional attributes ^16,39^. In some cases, it may be that functional attributes impacting host phenotypes diverge across closely related microbes ^40^, therefore the ASV level may be most appropriate. In other cases, a functional attribute selected for by the host may be conserved across wider evolutionary distances allowing for detection of h^2^ at higher taxonomic levels. Uncovering the appropriate unit for calculating heritable signal in host associated microbial communities will be an important challenge for future studies.

### Genetic architecture of host-microbiome interactions in roots

We identified numerous regions of the host genome associated with the abundance of core taxa. In addition, our results indicate that associated SNPs passing a genome wide threshold are rarely shared across multiple ASVs, yet the tails of GWAS p-value distributions contain commonly associated loci. These results suggest that loci with the largest effects on any particular ASV’s abundance are specific to that microbe while loci with smaller effects are shared between ASVs. Together, these results indicate that microbiome assembly is a complex trait given that the microbiome constitutes a consortium of interdependent bacteria; that many significant loci were identified associated with these microbes’ abundances; and that many GO term enrichments were uncovered associated with these loci. That is, many genes and processes contribute relatively small effects to influence the relative abundance for various ASVs.

A difficulty in presenting these data is their complexity and the plethora of uncovered candidate genes putatively involved in microbiota assembly. We therefore focused on loci impacting the most members of the microbiome (i.e. pleiotropic loci, Fig 4). Several compelling candidate genes were identified among the commonly associated loci which showed enriched expression in roots. Among these were a cellulose synthase subunit, whose ortholog in Arabidopsis is involved in secondary cell wall synthesis and has been reported to influence resistance to soil-borne bacterial pathogens in a defense hormone-independent manner ^41^. We also identified two root-expressed candidate nucleotide-binding leucine rich repeat proteins (NLRs) showing associations to multiple ASVs. NLRs are important sensors involved in effector triggered immunity and have been implicated in affecting sorghum rhizosphere microbiota ^17^. Given the diversity of NLR genes within plant species (switchgrass has well over 1000 annotated NLR genes) and the presence / absence variation between individuals within species ^42^, an open question is how the repertoire of NLR genes shapes root associated microbiota. The co-evolution between NLR genes and microbiota will remain an compelling hypothesis to explain local adaptation to the biotic environment and may serve as a means for fine-tuning microbial communities. Ultimately, uncovering specific mechanisms and genetic networks controlling microbiota assembly requires reverse genetic approaches. Several studies in maize have used mutants to show that ablation of specific metabolites in exudates can modify microbial community composition ^43^ and can lead to a significant impact on plant resistance to herbivory ^44^. Our study provides a list of possible candidate loci to target for future research.

### An association between Pattern-triggered immunity and root microbiota composition

Several of our analyses implicated physical and immune defenses as modulators of microbiome composition. In our study we investigated the role of plant genotype in explaining PTI variation using the elicitor flg22. While flg22 is one of many known elicitors, it serves as a good proxy for PTI given that pattern recognition receptors share similar co-receptors which funnel into similar pathways ^45^ and downstream transcriptional responses show strong overlaps ^46^. Much like a recent study in Arabidopsis, our results revealed strong heritable variation in PTI response within our population ^47^. Further, our analysis revealed a link between the abundance of the ATX core microbiota and modes of PTI variation within our switchgrass population. Particularly strong associations, both negative and positive, were observed between the first axis of PTI variation (ROS burst magnitude) and a phylogenetically broad set of root-associated microbes (Fig 6D). PTI canonically inhibits the entry of perceived pathogens ^48^, but our results suggest that it may also gate or limit the proliferation of commensal bacteria and their interactors, at least for ASVs with negative genetic covariances. This result is in line with previous studies showing that attenuation of PTI can lead to altered microbiota composition and even dysbiosis ^49^. Similarly, mutant Arabidopsis plants with altered defense hormone production host atypical root microbiota, indicating that immune signaling is an important modulator of microbiota assembly ^50^. On the other hand, we found ASVs with strong positive genetic covariance with PTI. These ASVs may 1) stimulate PTI sensitivity, such as in the case of induced systemic resistance (ISR); 2) escape the effects of PTI; or 3) benefit from the exclusion of PTI sensitive microbes. Deciphering the role and mechanisms of the host immune system in regulating microbiota assembly processes and how assembly of microbiota in turn modulates the host immune system is an active area of investigation with implications for the design of plant probiotics ^28^.

## CONCLUSION

We found that though environmental variation in natural field locations is the primary driver of microbial community composition, host genotype leaves a significant, widespread footprint on the root microbiome. We find evidence that locally adapted host genotypes enrich highly prevalent local microbes compared to foreign genotypes. Leveraging the associations with microbiota via manipulation of host genetics to favor desirable outcomes on plant fitness or yield is a goal that is currently unrealized. By characterizing which microbes are responsive to plant genotype and potential loci involved in host-microbiome interactions, the insights from this study may be of use when engineering or configuring associations between plants and microbes in the field.

## METHODS

### Plant collection, propagation, and planting

Collection, propagation, and field planting of the switchgrass population was previously described by Lovell et al. Briefly, the diversity population was established by collecting seeds and rhizomes from natural as well as common garden resources and transported to Austin, TX where the accessions were clonally propagated. Switchgrass is an outcrossing perennial plant, hence individuals in the planting populations are clonally propagated ramets and it is not possible to raise identical plants from seed. The genomes for individuals within the population were resequenced, aligned to the reference genome, and genomic variants were identified. Initial growth of plants and seedlings occurred in a mixture of Promix peat-based potting soil and calcined clay (Turface). Rhizome propagules were transplanted into 5 gallon pots containing finely ground pine-bark mulch and nutrients were supplied through slow release fertilizer (14-14-14, Osmocote). Final propagation of the accessions occurred in 2018 where ramets were grown in 1 gallon pots containing pine-bark mulch. In May to June 2018 the ramets were transplanted into the common gardens. Briefly, the fields were covered with weed cloth and the layout was marked such that each plant had a minimum of 1.56 m from the four surrounding plants. Holes were cut into the weed cloth and the soil was excavated using a spade shovel. The plants were placed into the holes, surrounded by soil, and hand watered. The lowland cultivar ‘Blackwell’ was planted around the edge of the field sites to account for border effects.

### Root Sample Collection and Processing

Samples were collected in the summer of 2019. Samples from ATX were collected in June, 2019 while CMO and KMI samples were collected in early August or 2019. The gap in sample collection timing between the sites was intentionally set to account for phenological differences in AP13, the reference genome accession, between locations. The size of our plantings as well as various characteristics of switchgrass plants presented several challenges during sampling. Switchgrass plants are obligately outcrossing therefore cannot be destructively sampled. Given that microbiomes can be dynamic, and can potentially respond to weather events, sampling of the fields had to occur within one day. Our plantings are large, and a team of samplers was employed to quickly collect root samples. A 1-inch diameter punch core was used for sample collection. Briefly, the corer was placed at the edge of the crown and rotated to be tangential to the crown. This allowed us to avoid the original potting soil directly underneath the crown where the original transplantation occurred and minimized the chance of capturing legacy microbiota from the pre-transplanted roots. The corers were pushed 10-15 cm below the surface at a 45-degree angle. The soil-bound roots were extracted from the instrument using a scoopula and placed into a plastic baggie. Between samples, the corer was cleaned of remaining soil using a paper towel, but no effort was made to sterilize the instrument between samples as ethanol cannot remove DNA and bleaching / washing the instruments was not feasible for conducting the sampling in a reasonable timeframe. Roots were encased by surrounding soil in the core, therefore the risk of cross contamination was negligible. After a row was completed, the sampler returned to a workstation and the baggies were organized and placed into a cooler with ice packs or wet ice.

The samples were processed the next day. Living roots from the baggies were picked using ethanol and flame sterilized forceps. Two or three 1-inch pieces of roots were placed into a 2 mL tube with 1 mL sterile PBS. Typical root samples contained both transport roots with attached absorptive roots. The roots were vortexed in PBS for 10 seconds then sterilely transferred to a new, clean tube with 1 mL PBS. Again the roots were again vortexed to remove soil adhering to the surface and the resulting dirty PBS was discarded. This process was repeated until the PBS solution was clear and no soil remained in the tube. The roots in the tubes were then frozen and stored at -80 degrees until DNA extraction took place.

### DNA Extraction

DNA was extracted from samples using a procedure similar to Bollman-Giolai et al. ^51^. Briefly, root samples are ground to a fine powder with two sterile steel beads in a 2 mL tube using a GenoGrinder for 30s at 1750 rpm. After grinding 0.25 g of garnet particles (Lysing Matrix A, BioSpec) were decanted into the tube and 540 uL of Buffer I (181 mM NaPO4, 121 mM Guanidinium Thiocyanate) was pipetted into each tube. The samples were briefly vortexed, and 60 uL of buffer II (150 mM NaCl, 4% SDS, 500 mM Tris pH 8) was added. The samples were then placed into the Genogrinder for 2 min at 1500 RPM to grind / lyse. The tubes were centrifuged at 10,000 g for 1 min to palette debris. The supernatant (500 uL) was transferred to a deepwell (1mL) 96-well plate and 250 uL of Buffer III (133 mM Ammonium Acetate) was added to the samples and vortexed to precipitate SDS and proteins. The plates were placed in 4 degrees for 5 min, then centrifuged at 4000 g. The supernatant (500 uL) was transferred to a new plate and 120 uL of Buffer IV (120 mM Aluminum Ammonium Sulfate Dodecahydrate) was added to precipitate fulvic and humic acids, typical PCR inhibitors from plant and soil samples. The samples were put at 4 degree for 5 min, then centrifuged for 2 min at 4000 g. After this step, the supernatant can be frozen /stored or directly used for the next SPRI bead purification step. For the SPRI cleanup, 300 uL of the supernatant is mixed with 240 uL of SPRI beads in a deepwell 96-well plate and incubated for 5 min. The plates were then placed on a magnet, allowed to clear, and the supernatant was discarded. The beads were then washed twice with 80% ethanol and allowed to dry for **5** min. DNA was then eluted using 50 uL of water and transferred to a 96 well plate for storage at -20.

### Library preparation and sequencing

We amplified the V4 region of 16S rRNA gene to survey microbial membership and relative abundance in the samples. We used a two-step strategy, where V4 regions were first amplified using modified primers published by Parada et al. ^52^. The primers were modified to add nextera sequencing primer annealing sites to the amplicons. The resulting PCRs were checked for amplification on a gel and cleaned using SPRI beads. The second round of PCR added barcodes and flow cell annealing adapters to the amplicons. Our barcoding strategy adds 12 bp Golay barcodes to both ends of the amplicon. The libraries were purified again using SPRI beads and quantified using Qubit high sensitivity assays. The amplicons were normalized for concentration by pooling samples at different volumes depending on their concentrations. The resulting pools were then concentrated using SPRI beads and run on a 2% agarose gel. The appropriate band was cut from the gel and purified (Nucleospin) and sent for sequencing.

Sequencing occurred at multiple centers. Our first two libraries were sent to both the HudsonAlpha Genomic Sequencing Facility and to the Joint Genome Institute (JGI). All of the other libraries were sent to JGI. All sequencing was performed using Illumina NovaSeq configured with the SP flowcell which is capable of 250 × 250 bp paired end read lengths.

### Sequence processing and ASV calling

Resulting reads were demultiplexed, if needed, using the demultiplex Python software (https://demultiplex.readthedocs.io/en/latest/index.html). Reads were trimmed to remove adapter sequences using cutadapt ^53^. ASVs were called using the dada2 R software package ^54^.

### Beta diversity measurements

Bray-Curtis dissimilarities were calculated using the *vegdist* function from the Vegan R package ^55^ on log2 transformed ASV relative abundances. Principal coordinate analysis was done using the *capscale* function from the Vegan package. Permanova was conducted using the *adonis* function.

### Modeling site and subpopulation effects on ASVs

We used a linear modeling framework to model the effect of field site, genetic subpopulation, and subpopulation x site effects on microbes. To be included in the analysis, an ASV must have been present in >= 50% of the total samples included in the study. For every ASV a linear model was run with the following structure

lm(ASV_abundancei ∼ log10(depth) + Site + Subpopulation + Site:Subpopulation)

Where ASV_abundancei is the vector of rank-based inverse normal transformation for the i^th^ ASV. This transformation was performed using the function RankNorm() from the R package RNOmni ^56^. Sequencing depth was accounted for by including the log10(depth) term in the model. Site represents the vector of field locations and Subpopulation represents the switchgrass genetic population of the host. Site:Subpopulation is the term capturing interaction effects between these two factors. Rank-based inverse normal transformations were performed to coax ASV relative abundances into a normal distribution, to fit the assumptions of the model. Variance partitioning of the terms was performed by running the function Anova() from the Car package on individual models and percent variance was calculated by dividing a factor’s sum of squares by the total sum of squares. Contrasts across model variables were calculated using the emmeans package ^57^.

### Genetic variance component analyses

Additive genetic variance and GxE variance was first calculated using the compound symmetry model in the R package Sommer. The compound symmetry structure model assumes constant total variance within each site as well as constant covariance between sites. This is the simplest model structure and was selected as the first step in our analysis because the model returns components for additive genetic variance and genotype by environment variance. To be included in the analysis, a feature must have been detected in >= 80% of the samples. The full model was run with the following structure.

Full_model <-mmer(rst ∼ Site + log10(depth), random =∼ vs(PLANT_ID, Gu=K) + vs(Site:PLANT_ID, Gu=EK), rcov = ∼units, data = x2, tolparinv = 1e-01, verbose = T)

rst is the vector of rank-based inverse normal transformed ASV relative abundance (or aggregated relative abundance if classification is above ASV). Rank-based inverse normal transformations were applied to the counts within each site for each ASV and resulted in a constant overall variance, fulfilling this assumption of the compound symmetry structure. In this model Site and sequencing depth were fit as fixed effects. PLANT_ID is the plant accession name and K is the kinship matrix with pairwise relationships between individuals in the population based upon SNP data. Site is the field location and ‘vs(Site:PLANT_ID,

Gu=EK)’ captures the variance of GxE in the model, where EK is a list of site-specific kinship matrices. Reduced models were constructed to test the contribution of VGxE and VA to the models. They were encoded as follows

reduced_1 <-mmer(rst ∼ Site + log10(depth), random =∼ vs(PLANT_ID, Gu=K), rcov = ∼units, data = x2, tolparinv = 1e-01, verbose = T)

Notably, this model lacks the GxE term ‘vs(Site:PLANT_ID, Gu=EK)’. This model was compared to the full model using a likelihood ratio test to examine whether GxE influenced the abundance of the tested ASV. To test for the effect of host genotype, we compared reduced_1 to the below model.

reduced_2 <-mmer(rst ∼ Site + log10(depth), rcov = ∼units, data = x2, tolparinv = 1e-01, verbose = T)

This model lacks the effect of genotype altogether, thus comparing reduced_2 to reduced_1 using a likelihood ratio test examining whether host genotype contributes to the observed variance of the tested ASV. To make a call on whether GxE or VA influenced microbial abundances, we first asked if GxE showed an adjusted P value < 0.1. If so, our analysis stopped and we flagged the tested ASV as showing significant GxE. If not, then we tested whether VA had an effect with an adjusted P value < 0.1. If so, we made a call that the ASV is affected by host additive genetic variance. If not, we inferred that the ASV was not affected by host genotype.

We next used the unstructured model in the sommer package to ask about additive genetic variance within each site. The unstructured model allows for unequal additive genetic variances within sites as well as unequal covariances between sites. This allowed us to ask about the influence of host genotype within sites and whether the influence of host genotype is consistent across multiple sites.

Multiple testing was accounted for through correction by the Benjamini-Hochberg approach, and a significant contribution of either parameter was determined at FDR < 0.1.

### Microbial Genome Wide Associations

We performed GWAS for microbes found in >80% of the samples within each site. For this analysis, where we were performing quantitative models, we removed samples where the focal ASV was not detected and the relative abundance were transformed as previously mentioned using the rank-based inverse normal transformation. GWAS was run using the SwitchgrassGWAS R package (https://github.com/Alice-MacQueen/switchgrassGWAS) ^22^. This package dynamically chooses the number of genetic PCs to include as covariates in the model to control for population structure and reduce genomic inflation. The SNP matrix used in the analysis was dense, composed of over 25 million SNPs generated from the *Panicum virgatum* V5 genome. The gene content near SNPs passing a threshold of 5×10^−8^ was generated using BEDTools window ^58^ on the *P. virgatum* v5.1 genome annotation with a window size of 50 kb.

For the core microbiota, i.e. microbes detected in >= 80% of the samples in each field site, the P-values for the GWAS scans of each microbe were combined using Fisher’s Method from the R package ‘metap’ ^59^.

### Detection of pleiotropic loci affecting multiple microbes

To identify regions of the host genome putatively influencing the abundance of multiple microbes we divided the genome into 25 kb bins, consistent with average linkage equilibrium decays suggested in other switchgrass studies ^60^. For each microbe, this resulted in 43,402 bins. We next calculated the minimum p-value of the SNPs within each bin for each microbe and retained the top 0.5% of bins with the lowest p-values (217 bins). The resulting QTL bins were then compiled into a presence / absence matrix and we kept the top 5 loci from each site for further analysis. We tested the likelihood of observing the number of overlapping loci in our data by using a permutation framework. In our QTL x ASV matrix, the ASVs were the rows and QTL were the columns. We randomized the QTLs for each ASV in the matrix and counted the maximum number of overlaps, stratifying by field location. This was performed 1000 times to develop a null distribution. All of our top 5 pleiotropic loci had p < 0.001. We chose to only analyze the top 5 loci for each site for presentability, but include the other loci passing this significance threshold in the supplemental tables.

### Gene Ontology Enrichments

We identified the gene content of the QTL matrix composed above using bedtools window, then extracted the Gene Ontology categories for each gene within each 25kb genomic bin. Enrichment was calculated against the background genome GO counts using a hypergeometric test and P values were corrected for multiple tests using the Benjamini-Hochberg procedure.

### Gene Expression Analysis

The expression values for gene underlying putative pleiotropic loci were extracted from the *Panicum virgatum* gene expression atlas which can be found on Phytozome 13. The FPKM values for P. virgatum gene expression across tissues and environments were generously shared with us by the group of Jeremy Schmutz. Differential expression between root and shoot tissue was performed using the following linear model on FPKM values.

lm(log2(expression) ∼ Tissue)

The resulting P-values for the term ‘Tissue’ were corrected using the Benjamini-Hochberg procedure and significance was called at adjusted p value < 0.05.

### Pattern Triggered Immunity Assays

Leaf tissue was collected from the ATX field site plants in the spring of 2020. Leaf disks were punched from the leaves on location in the field and immediately placed in 2 mL of sterile DI water in a 48 well plate and covered with aluminum foil. The plates were gently shaken for 2 hours, then the disks were transferred to white, opaque 96 well plates in 50 uL of sterile DI water, wrapped in aluminum foil, and left overnight. The next day, the disks were treated with 50 uL of Flg22 elicitor cocktail (10ug/mL horseradish peroxidase, 34 ug/mL L-012, and 1 uM Flg22). The plates were read over a time series on a SpectraMax M3 plate reader. Negative control plates with a randomly selected group of genotypes were mock treated (10ug/mL horseradish peroxidase, 34 ug/mL L-012, water). Each genotype was read in triplicate. To analyze the data, we log transformed the relative luminescence units of the time series and reduced the dimensionality using PCA.

### Genetic covariances of PTI axes and bacterial abundances

We performed genetic covariances between the first three PTI PCA axes and ATX root microbe relative abundances using the R package Sommer. We used the following mixed effects model.

covar_mod <-mmer(cbind(ASV_abund, PTI_PC) ∼ 1, random= ∼vs(PLANT_ID, Gu=K), data=data, tolparinv = 1e-1)

The terms for ASV_abund and PTI_PC changed depending on the focal ASV and focal PTI PC axis. Covariance estimates and standard errors for the estimates were gathered using the following command.

covar <-vpredict(covar_mod, covar ∼ V2 / sqrt(V1*V3))

P values for observing the covariance estimate or larger (in magnitude) were calculated as p = 2*pnorm(estimate / standard_error, lower.tail=FALSE)

## Figure Legends

*Supplementary Figure 1. Sequencing depths for samples included in this study*

*Supplementary Figure 2. Comparison of the results from the compound symmetry and unstructured models used to estimate genetic variance components contributing to the abundance of ASVs. How ASVs change in their assignment of significant Va (G), GxE, or no association to host genetic variation (y-axis) between the two model structures (x-axis) are denoted by lines. The number of ASVs changing assignments are denoted by line thickness and written values*.

*Supplemental Figure 3. Covariances of the same ASVs compared across different sites*. A) Density plots showing the distribution of covariance estimates. B) Number of ASVs with significant covariance.

*Supplemental Figure 4. GWAS reveals loci contributing to community structure in each field site*. GWAS on the first three PCo of community dissimilarity metrics (Bray) from each field location. The genome-wide threshold, set at 5×10^−8^, is indicated by a dashed line in each Manhattan plot.

*Supplemental Figure 5. ASV by site GWAS scans identify diverse taxa affected by genomic variation*. Bacterial ASVs tested for and showing significant associations with SNPs (P < 5×10^−8^) in A) Austin, TX, B) Columbia, MO, and C) KBS, MI. The number of tested microbes is in black while ASVs with significant associations show up in the color corresponding to the field site. The inset in panel C is the association between h2 and having at least one SNP associated with microbial abundance. D) Heatmap of ASVs where GWAS was performed in multiple sites. Black boxes indicate microbes with at least one significant SNP associated with relative abundance.

*Supplemental Figure 6. Gene Ontology enrichments show similar and contrasting patterns across locations*.

*Supplemental Table 1* Study-wide and site-specific core taxa

*Supplemental Table 2* Subpopulation specific enriched microbes

*Supplemental Table 3* Compound Symmetry Model Results

*Supplemental Table 4* VA estimates using unstructured model

*Supplemental Table 5* Statistical Pleiotropic Loci

*Supplemental Table 6* Proportion of microbes with enriched GO terms

*Supplemental Table 7* Enriched GO terms from GWAS meta-analysis

*Supplemental Table* 8 Significant GWAS Metanalysis Annotations

## ACKNOWLEDGMENTS

This research was supported by the Office of Science (BER), U.S. Department of Energy, Grant no DE-SC0014156 and DE-SC0021126. The work (proposal: 10.46936/10.25585/60000507) conducted by the U.S. Department of Energy Joint Genome Institute (https://ror.org/04xm1d337), a DOE Office of Science User Facility, is supported by the Office of Science of The U.S. Department of Energy operated under Contract No. DE-AC02-05CH11231. This work was supported in part by the Great Lakes Bioenergy Research Center, U.S. Department of Energy, Office of Science, Office of Biological and Environmental Research under Award Number DE-SC0018409. Support for this research was provided by the National Science Foundation Long-term Ecological Research Program (DEB 1832042) at the Kellogg Biological Station and by Michigan State University AgBioResearch. J.E. acknowledges the support of the USDA National Institute of Food and Agriculture Postdoctoral Fellowship (grant no. 2019-67012-2971/project accession no. 1019437). In addition, we would like to acknowledge Allison Hutt, Nick Ryan, and Lisa Vormwald for their help in collecting samples.

