## Supplementary figures and images for "Heritability and host genomic determinants of switchgrass root-associated microbiota in field sites spanning its natural range"

### Supplemental Figures

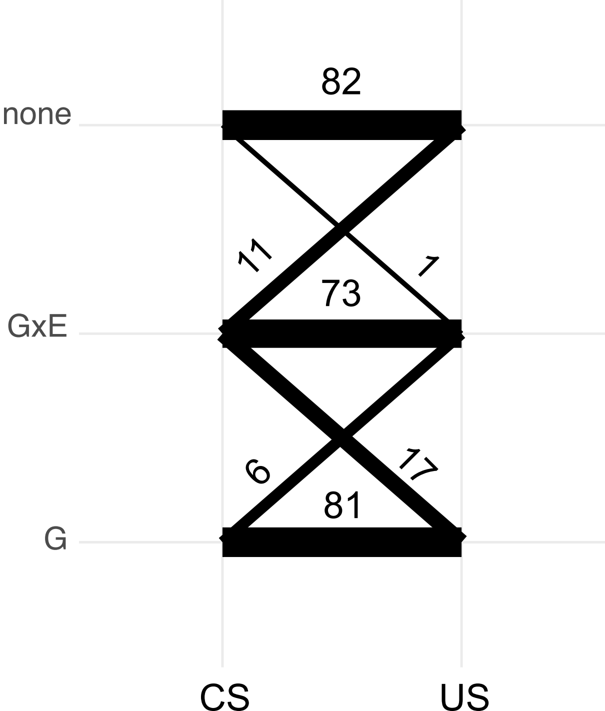

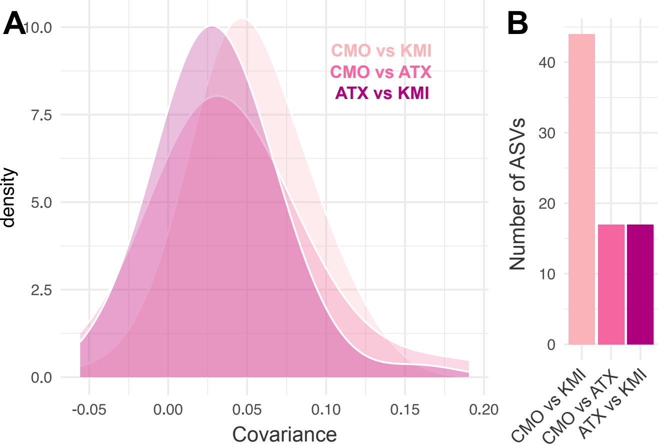

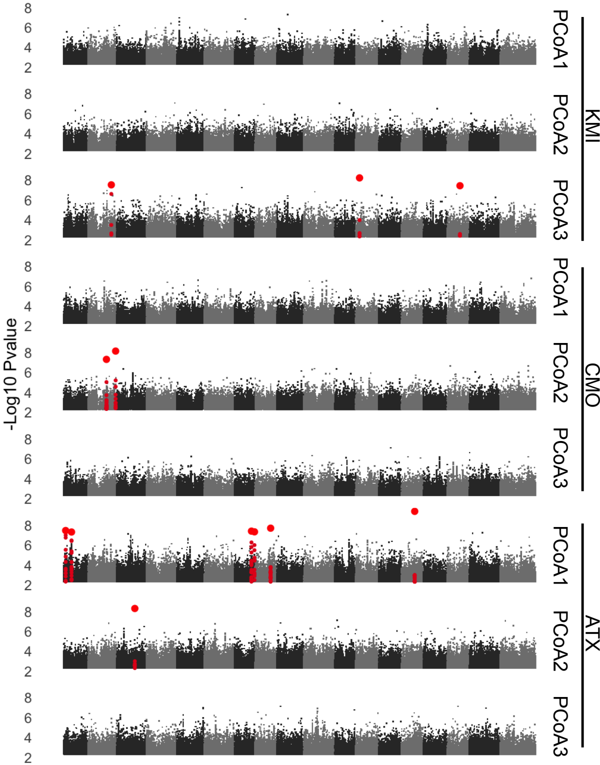

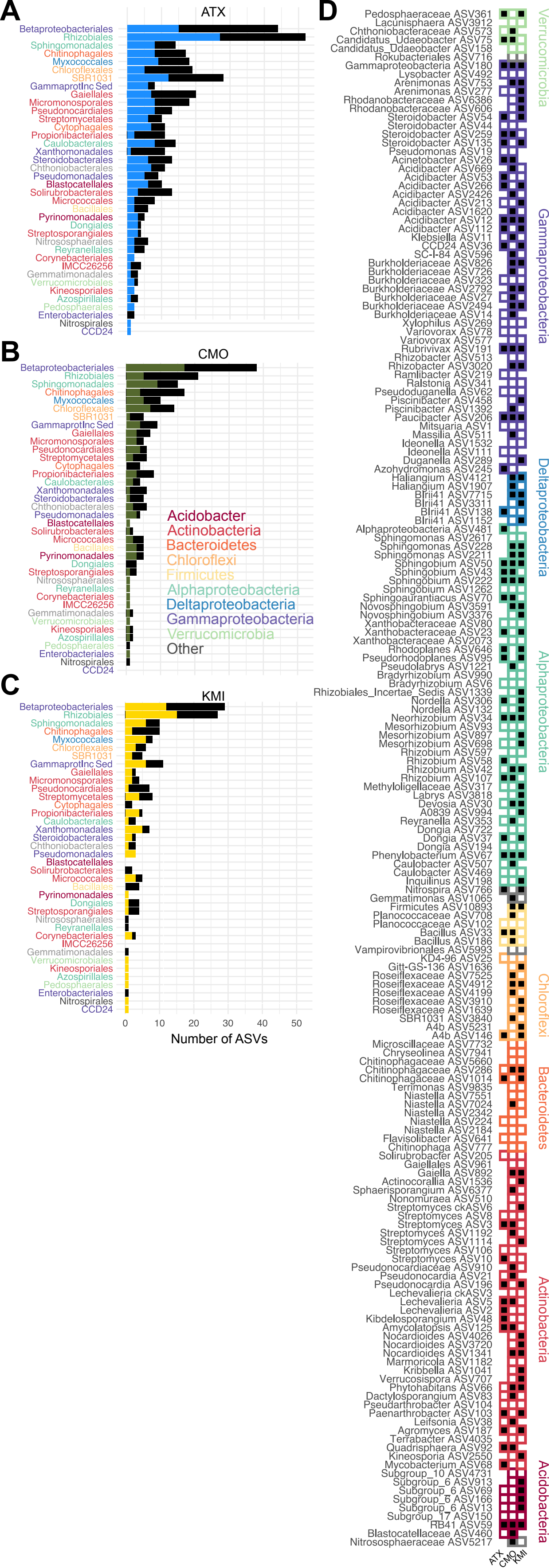

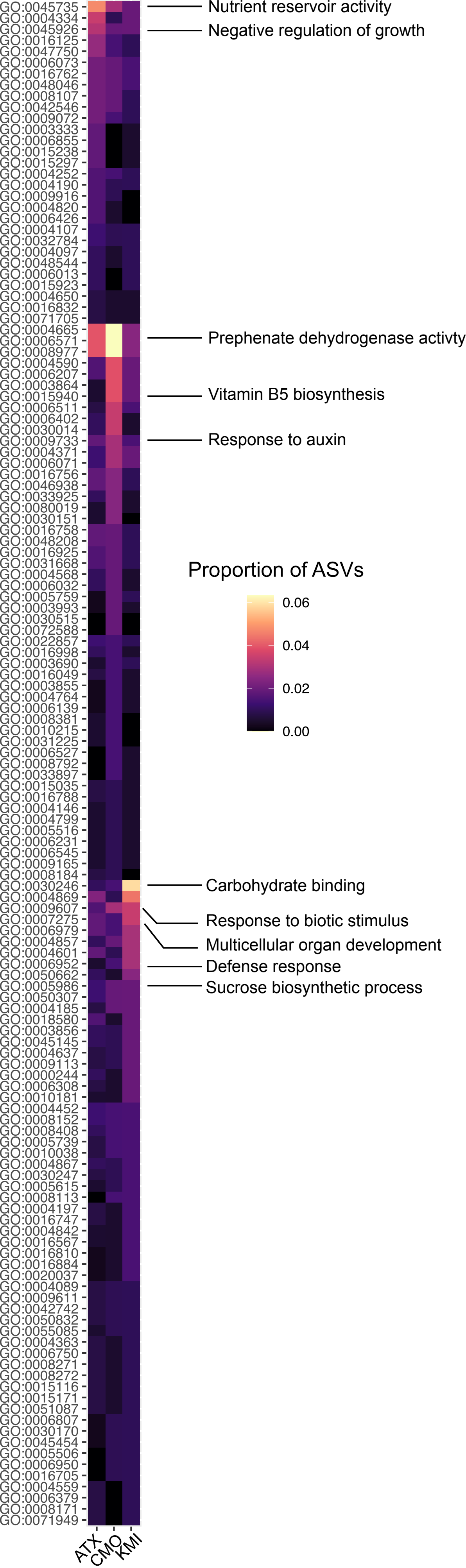
